# Single haplotype admixture models using large scale HLA genotype frequencies to reproduce human admixture

**DOI:** 10.1101/336693

**Authors:** Alexandra Litinsky Simanovsky, Abeer Madbouly, Michael Halagan, Martin Maiers, Yoram Louzoun

**Affiliations:** Department of Mathematics and Gonda brain research institute, Bar-Ilan University, Ramat-Gan 52900, Israel; Bioinformatics Research, Center for International Blood and Marrow Transplant Research, Minneapolis, MN, USA

**Keywords:** HLA, genetic admixture, non-negative matrix factorization, stem-cell donor registry

## Abstract

The Human Leukocyte Antigen (HLA) is the most polymorphic region in humans. Anthropologists use HLA to trace populations’ migration and evolution. However, recent admixture between populations masks the ancestral haplotype frequency distribution.

We present an HLA-based method based on high-resolution HLA haplotype frequencies to resolve population admixture using a non-negative matrix factorization formalism and validated using haplotype frequencies from 56 populations. The result is a minimal set of original populations decoding roughly 90% of the total variance in the studied admixtures. These original populations agree with the geographical distribution, phylogenies and recent admixture events of the studied groups.

With the growing population of multi-ethnic individuals, the matching process for stem-cell and solid organ transplants is becoming more challenging. The presented algorithm provides a framework that facilitates the breakdown of highly admixed populations into original groups, which can be used to better match the rapidly growing population of multi-ethnic individuals worldwide.

**Author Summary:** Human Leukocyte Antigen (HLA) is known to be the most polymorphic region in the human genome. Anthropologists frequently use HLA to trace migration and evolution of different populations. This is due to the high linkage among HLA genes leading to the transmission of intact haplotypes from parents to offspring, hence preserving key population ancestral features.

We developed a new HLA-based method to identify admixture models in mixed populations using high-resolution HLA haplotype frequencies. Our results highlight that a single highly polymorphic locus can contain enough information to map clearly human admixture and the population genetics of the different human populations, and reproduces results based on SNP arrays.

The presented algorithm is validated using haplotype frequencies sampled from 56 worldwide populations. Under such factorization we demonstrate that 90% of the variance in these populations can be explained using a much-reduced set of 8 ethnic groups. We demonstrate that the estimated ethnic groups and admixture models agree with the geographical distribution, population phylogenies and recent historic admixture events of the studied populations.

## Introduction

Human Leukocyte Antigen (HLA) plays a crucial role in both the adaptive and innate immune system and is proven instrumental in multiple medical disciplines, including matching for solid organ and hematopoietic stem-cell transplantation (HSCT) [1-4]. As generations of humans migrated throughout the world, HLA has evolved with each population conserving key ancestral features [5], and with more than 15,000 defined alleles to date[6] and over 1,000,000 haplotypes [7], it stands as the most polymorphic region in the human genome.

The study of human admixture and population stratification addresses a growing need in multiple disciplines, including large-scale disease and drug association studies [8],[9], planning and modeling volunteer adult donor stem-cell registries [10] and personalized and consumer genetics [11]. Typical population genetics algorithms interrogate Single-Nucleotide Polymorphisms (SNPs) in large genomic or genome-wide regions to track concepts like population size, migration, selection, admixture, recombination or genetic drift[12-18] or study the population genetics of specific ethnic groups [17, 19, 20]. The need for large genomic regions stems from the limited polymorphism in bi-allelic SNPs. This, however, is not the case for HLA. With the evolution of HLA over millions of years, distinctive features of the MHC locus became evident, such as gene density, diversity, and low recombination [21, 22]. Multiple models have been proposed to explain HLA polymorphism [23, 24], the most prominent being pathogen-driven balancing selection [25]. This extensive polymorphism facilitates tracking human migration patterns, population admixture, allelic diversity, pathogen evolution and selection through the study of HLA allele and haplotype frequencies [26]^-^[27].

Importantly, because of the high-gene density, diversity and low recombination rates, HLA alleles are inherited as haplotypes, preserving a high degree of linkage disequilibrium (LD) [28]. Recent evidence shows a clear purifying selection in HLA haplotypes [29], i.e. many combinations of HLA alleles are seldom separated over multiple generations. One interpretation is that certain allele groupings on a haplotype are tuned to work together. Hence, the absence of detectable recombination may reflect maintenance of haplotypes with favored immunological function through selection against unfavorable recombinant haplotypes[21], creating founder haplotypes with insufficient evolutionary time to degrade. The high occurrence of a handful of distinct long-range HLA haplotypes has long been of interest because of their prevalence and strong associations with complex diseases [30, 31] [32].

Multiple researchers have studied population genetic traits through the estimation and analysis of HLA allele and haplotype frequencies. Klitz et al [33] studied detailed Expectation Maximization based on three-locus low resolution haplotype frequencies to estimate the phylogeny of Jewish populations using shared haplotypes and correlations between haplotype frequencies among populations. More detailed correlation based analysis of high resolution six-locus haplotype frequencies was performed by Gragert et al [34] where population groups and phylogeny was detected among the main US populations as defined in the Be The Match^®^ US volunteer donor stem-cell registry. Recently, we studied the selection affecting HLA haplotypes [29, 35]. However, to our knowledge there is no published work dichotomizing HLA haplotypes into original populations that break down the admixture in haplotype distributions.

Population substructure is often estimated using admixture models, where each population is broken down into a mixture of original populations [36] [37]. This is formally a hidden state model, where the hidden states and the probability distribution of each hidden state in each population are estimated. To resolve the admixture into its primary components, most algorithms use a maximum likelihood approach [37, 38]. Multiple methods have been developed to detect population admixture (e.g. ADMIXTURE [37] and STRUCTURE[36, 39]). However, hidden state model methods can produce spurious results in the presence of background LD, which often happens in admixed populations due to multiple effects, such as genetic drifts and in some polymorphic markers, such as HLA and microsatellites [40]. Additionally, these models are not adapted to a single highly polymorphic region in contrast to large regions of limited polymorphism. We here present a method to estimate population admixture, in the presence of background LD, based on Non-Negative-Matrix Factorization (NMF). Using HLA haplotype frequencies from 56 world populations, we decompose the frequency distributions into the non-negative composition of a small number of original populations (OP) and test the precision of our results using a large dataset of high-resolution HLA haplotype frequencies, estimated from over 3.5 million individuals. Data were collected from volunteer unrelated adult stem-cell donor registries worldwide for haplotype frequency analysis and registry modeling studies. We then show that these original populations can be used to detect the admixture composition of new populations.

## Materials and Methods

### Study Populations

HLA haplotype frequencies from 56 global populations were used in this study (table 1-a). These populations were predominantly volunteer donors in stem-cell donor registries across the globe. We used haplotype frequencies from the Ezer-Mizion Registry in Israel [41] multiple registries in India[42], the US Be The Match registry® [34], a number of European registries that list their donors within the US registry (The Netherlands, Norway, Sweden, Wales) and the Australian registry, including two registries that list within it: New Zealand and Thailand. Haplotype frequencies from the Canadian registry were used for method validation (table 1-b[34]). Five-locus HLA-A∼C∼B∼DRB1∼DQB1 allele-level haplotype frequencies were used for all populations except for the Canadian groups for which only four-locus HLA-A∼C∼B∼DRB1 frequencies were available.

**Table 1.**
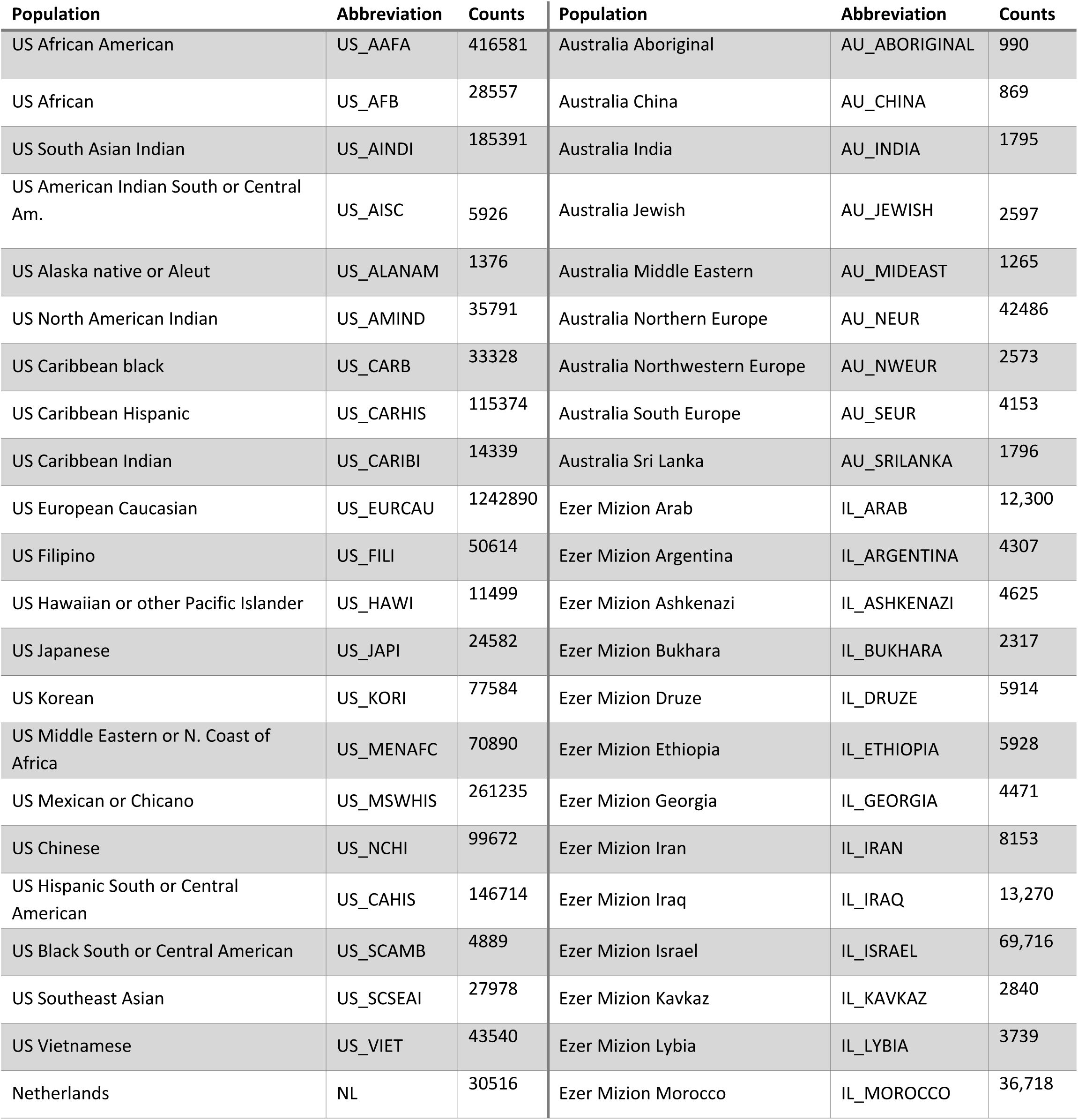
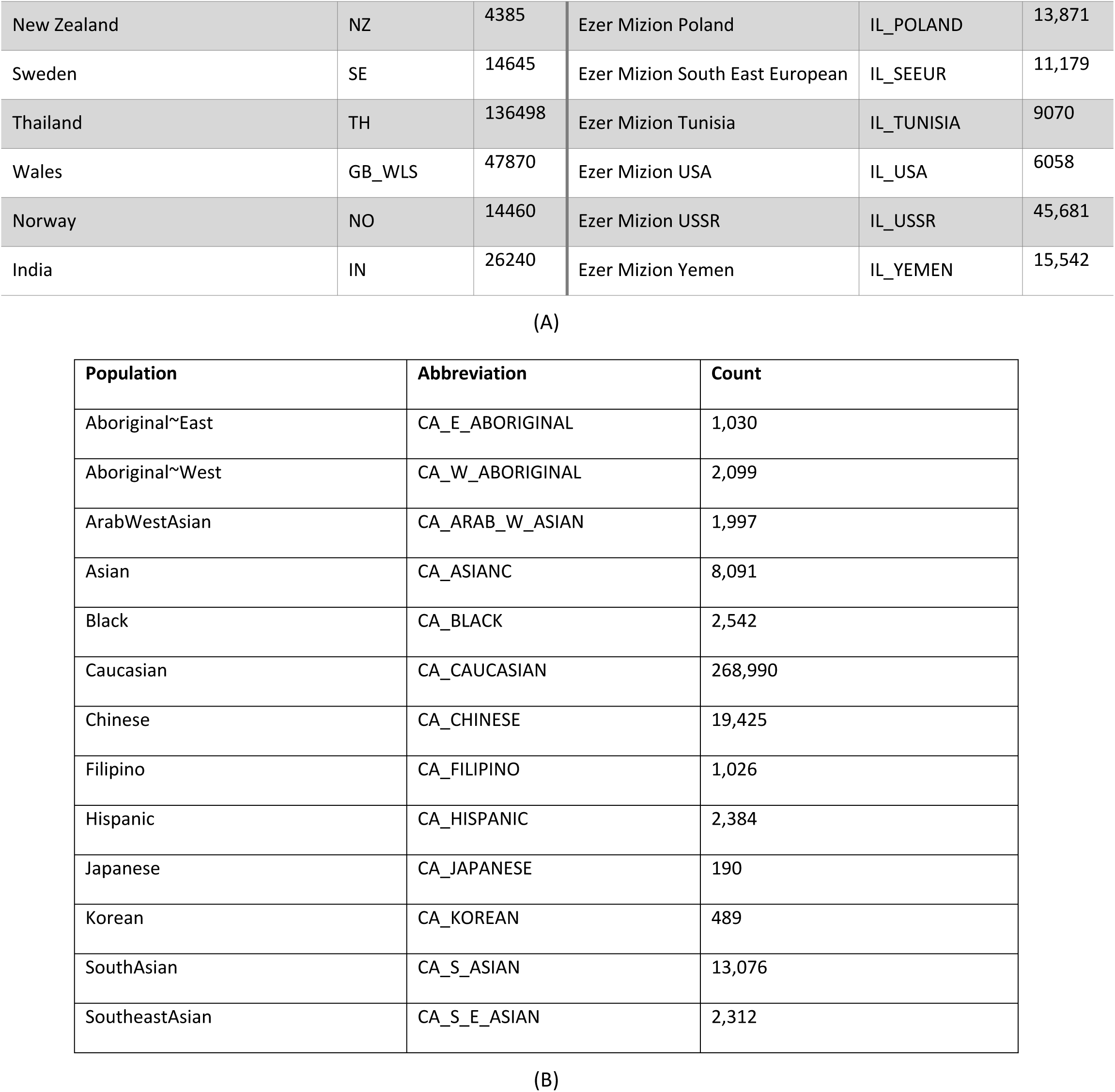
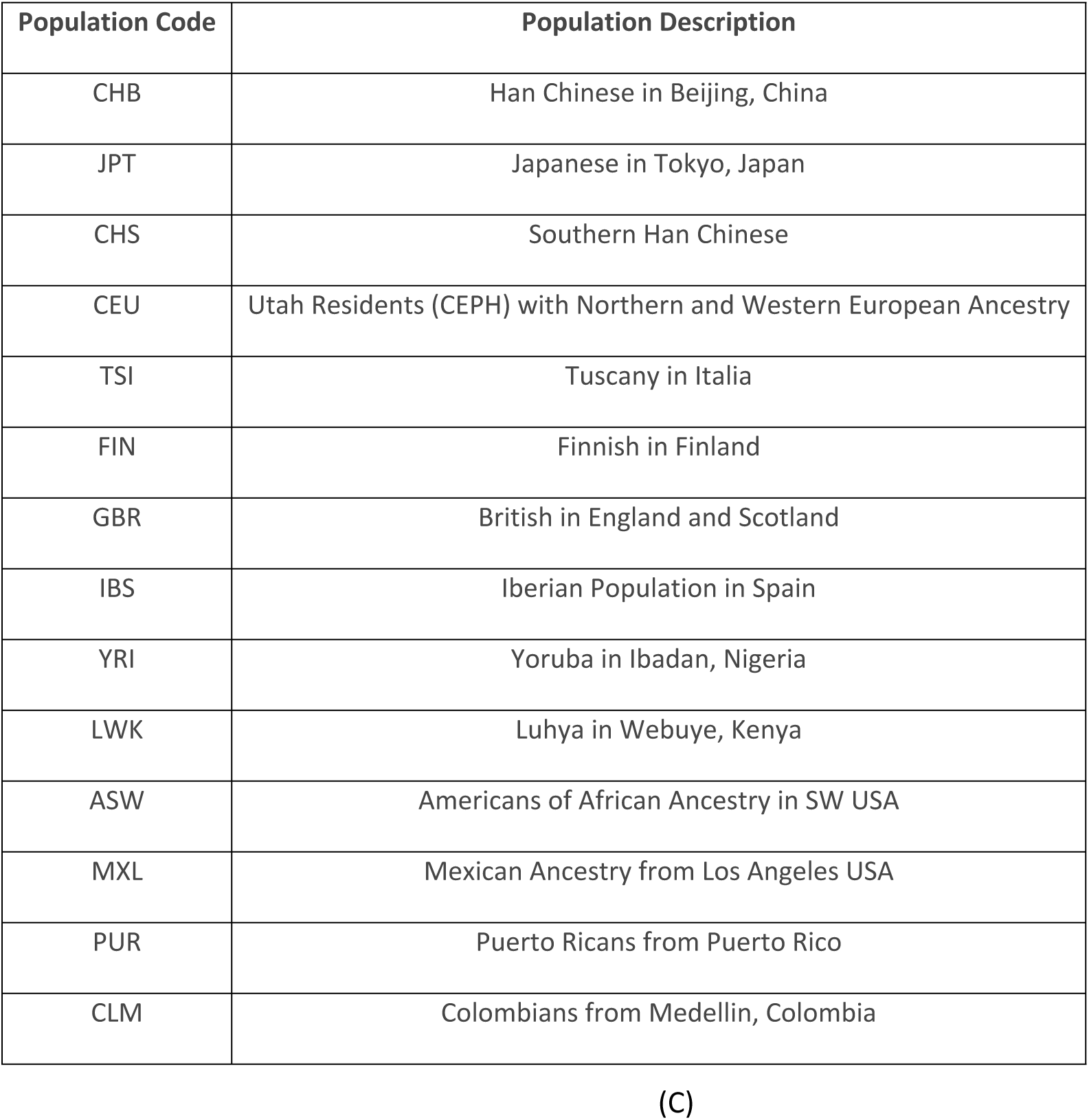
A) Study populations, acroynyms used and the sample size used to generate haplotype frequencies for each population. B) Populations from the Canadian OneMatch registry used for validation. C) Phase I 1000 Genome population acronym details

### Nonnegative Matrix Factorization

To estimate population admixture proportions using Nonnegative Matrix Factorization (NMF), a non-negative matrix **C** would be factorized into two non-negative matrices **A** and **B**, where **C** is a ***n****X****m*** matrix, representing the haplotype frequencies in ***n*** populations, each with ***m*** haplotypes. **A** is ***n****X****r*** non-negative mixing matrix and **B** is ***r****X****m*** non-negative “original” populations (OP) matrix. Each ***n***-dimensional vector represents one out of ***m*** populations. In the current application, ***m*** constitutes the 56 studied populations and ***n*** is of the order of a million different haplotypes. The variable ***r*** is chosen to be much smaller than ***n*** and ***m***. The range of **r** values, representing the count of possible OPs, was tested between 1 and 23. The results of the decomposition are a function of the chosen cost function [43]. We have tested the Frobenius, Kullback-Leibler (KL), Lee, offset and LS-NMF(least squares non-negative matrix factorization) cost function as implemented in the R package NMF [44].

The factorization is initialized with a seed (e.g. A_0_ and/or B_0_), which initiates the expectation maximization (EM) process. In each iteration, a new non-negative matrix A or B is calculated. Then, the new obtained matrix is used for calculating the complementary matrix. Iterations are repeated until converging to a locally optimal matrix factorization [43].

There are currently multiple standard algorithms for NMF [43]. These algorithms approximate a non-negative matrix (a matrix where all elements are non-negative) as a product of two low-rank non-negative matrices. The results of the approximation are affected by the loss function used and by the weighting of the input columns. In the current context, we produce a haplotype frequency matrix of all HLA haplotype frequencies for over 50 populations and present it as the product of eight original population frequency matrix and mixing factors.

### Phylogenetic analysis

We applied a Neighbor-Joining (NJ) algorithm to construct a phylogenetic tree of the studied populations [45, 46]. Following the tree construction, populations were grouped by the most dominant OP obtained from the NMF analysis. To validate the NMF algorithm, we investigated the similarity between the tree structure and the phylogenetic grouping.

The NJ distance matrix was based on the genetic distance between the population haplotype frequencies. We used the pairwise Fst measure *[47, 48]* calculated for all combinations of population pairs. Typically, the Fst value ranges from 0.0 to 1.0, where 0.0 indicates identical population frequencies and 1.0 indicates significantly different populations. We use the Fst algorithm as implemented by Weir et al*[49-52]*.

Principal Coordinate Analysis

Principal Coordinate Analysis (PCOA) (also denoted multi-dimensional scaling) was applied to translate the distance matrix into a 3-dimensional Euclidean space. The projection was performed using the wcmdscale function implemented in the R package, Vegan.

### Hidden state model algorithm

We compared the performance of our NMF-based admixture algorithm to the software STRUCTURE[36] that estimates population substructure using a Markov Chain Monte Carlo (MCMC) Model. Given that STRUCTURE produces spurious results for markers with excessive background LD (such as HLA), we ran STRUCTURE on select SNP genotypes from the reference 1000 Genomes phase I dataset [53] and compared the output admixtures to admixtures estimated by OPs in populations of similar ancestry as the 1000 Genome populations in our study. The 500 selected genome-wide SNP ancestry informative marker panel was previously published and validated for delineation of population substructure [54]. STRUCTURE v2.3.4 was run using K = 8 clusters (like the count of OPs), with 7,000 replicates and 50,000 burnin cycles.

### Inference of unlabeled populations

A possible application of the work presented here is to dissect an unknown admixed population into a set of OPs. Starting with a set of estimated OP haplotype frequencies, one can decompose a new population of unknown admixture using a regression of these frequencies over the used original frequencies.

Often the new population may have lower resolution (e.g. 4 locus information instead of 5). In such a case, the original frequencies are reduced to a similar resolution prior to regression.

## Model

We present a model to infer the admixture of a population using haplotype frequency distributions from multiple populations. The 56 studied populations originate from multiple ethnic groups in four continents with ancestral roots encompassing all six world continents. Some populations were relatively homogeneous (e.g. Vietnamese) while others were formed of relatively recent admixtures (e.g. African-American and Latin American)[54, 55]. To estimate the components of a population admixture, we assume the existence of K OPs, each represented by a haplotype frequency distribution denoted as *P*_*j*_, where *j* ranges from 1 to K OPs. Each observed population is assumed to be a positive normalized combination of such OPs:

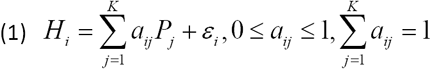

where *ε*_*i*_ is a noise vector that is affected by the sample size and may vary for each haplotype. To translate (1) into a matrix decomposition problem, we rewrite it as:

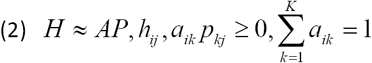

where A is the matrix of all *a*_*ij*_. To estimate the number of OPs best describing the studied populations, an appropriate cost function is required. Such a function can be based on the error in the reconstructed populations (*E* = *H* – *AP*) and is scaled by the sample size of each population. Such scaling may be crucial, since some populations are over-represented in the study cohort. Moreover, if the population is not scaled, splitting a population into two sub-population will double its contribution to the error. Other cost functions were also tested (See Supp. Mat. Table S1 for a list of tested cost functions and weightings). The relative error of the decomposition was computed for each algorithm as a function of the number of used OPs:

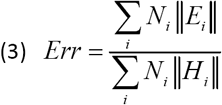

Where *E*_*i*_ and *H*_*i*_are column vectors of E and H respectively representing a single population, *N*_*i*_ is the population size, and the norm used is Euclidean. In all tested cost functions, the errors plateaued at 7-10 OPs (Fig. 1A). Note that non-scaled solutions produce OPs strongly biased toward highly admixed groups (Supp. Mat. Fig. S1) and produce a higher error rates (Fig. 1A).

**Figure 1:**
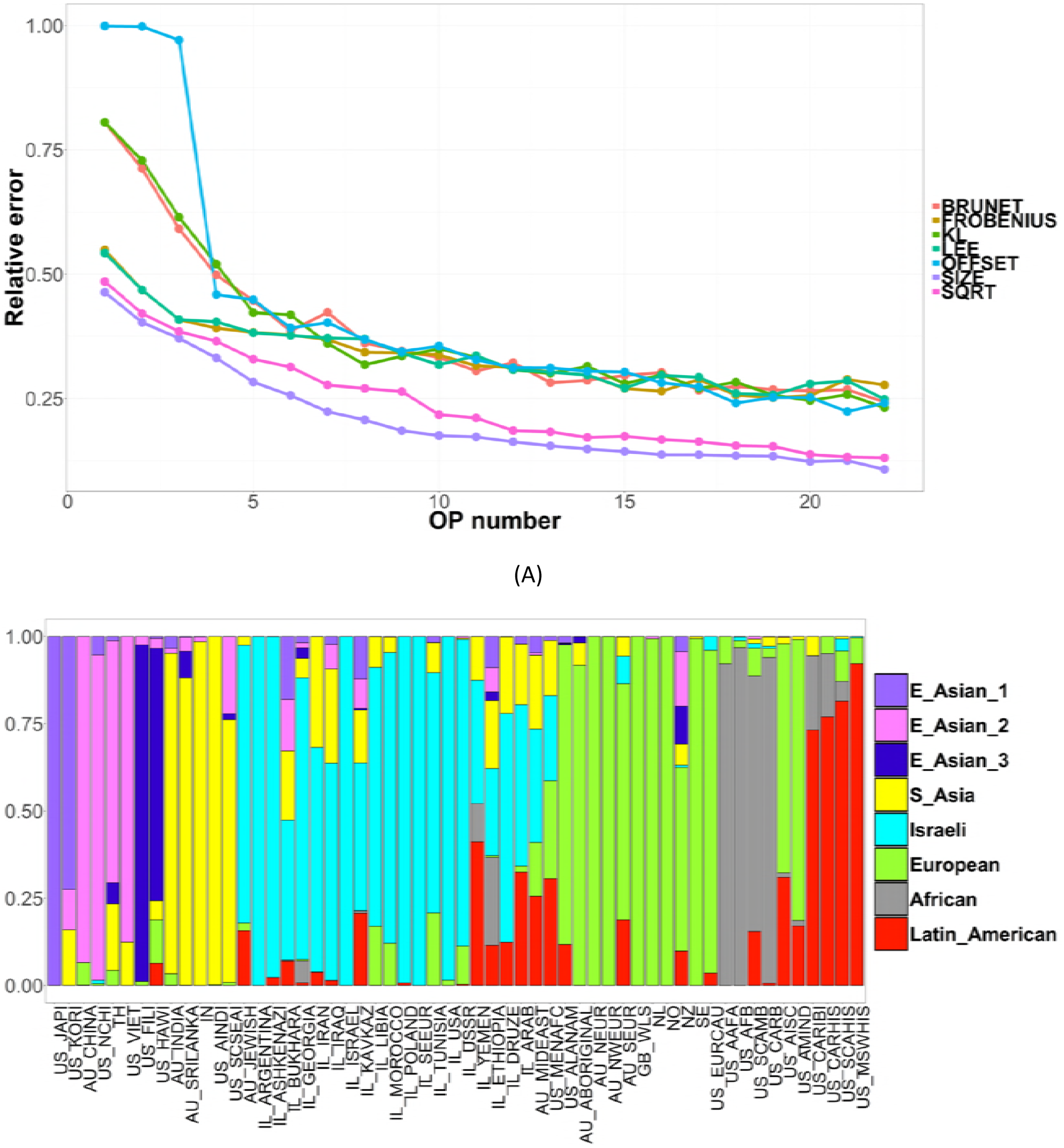
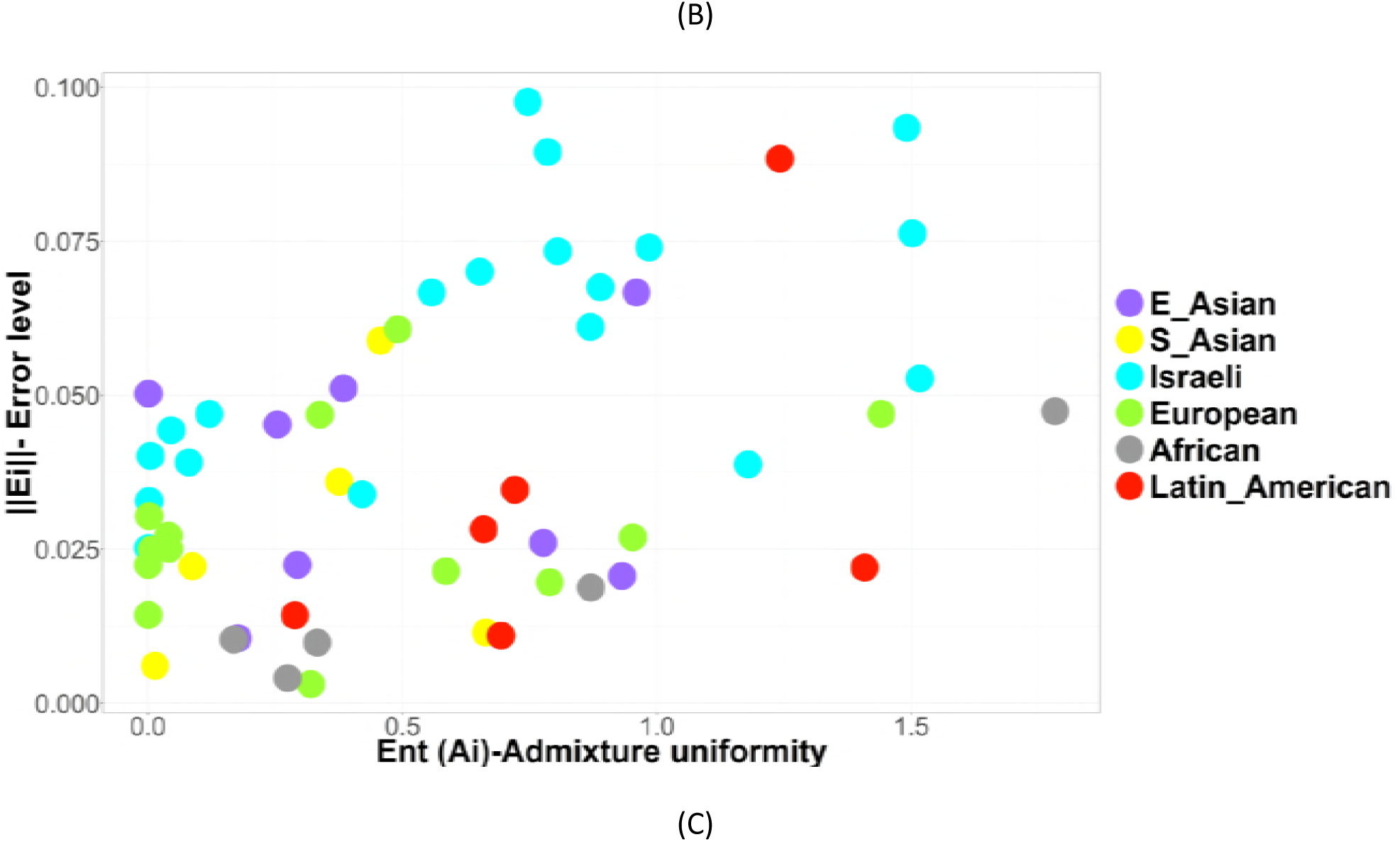
A) Relative decomposition error computed for each evaluated cost function as a function of the number of OPs (Original Population). In all tested cost functions, the errors plateaued at 7-10 OPs. B) Population Admixture Analysis: individual ancestry proportion of 56 population samples with 8 OPs. The color composition reflects the admixture (i.e. the fraction of a given population explained by the appropriate OP. Some of the population are composed of a single OP. All population acronyms are listed in table 1. C) Scatter plot of admixture error ‖*E*_*i*_‖ vs. the admixture entropy *Ent*(*A*_*i*_). Each population has an assorted color, depending on the group it belongs to. The correlation between the entropy and admixture is unaffected by the population size.

Each OP is assigned a name according to the principal observed populations containing it, with the following main groups: European, African, Israeli, Latin American, South Asian and East Asian. When more OPs are used, the result is typically the splitting of one of these groups into sub-groups (Supp. Mat. Fig. S1). Given, the results of the admixture, we assign each observed population one main group according to the main OP composing them. In the following results, the observed populations are grouped according to the OP composing the largest fraction of the admixture vector of A) *A*_*i*_ (A single column of A) (Fig. 1B).

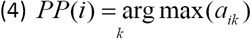

In all plots, populations are colored according to the OP with the *PP*(*i*) most contributing to this population.

## Results

Our results show that eight OPs are sufficient to represent the admixture in the 56 studied populations. We observed that the diversity level varied widely among populations; in some groups the admixture was predominantly represented by one OP as in the case of the US Japanese and Filipino populations, while other populations, such as US Middle Eastern, New Zealand and Australian Aboriginal were quite admixed (Fig. 1B). These results are consistent when the admixture method and number of OPs are varied (Supp. Mat. Fig. S1). The difference in the composition can be observed in the entropy of vector *A*_*i*_ -*Ent*(*A*_*i*_) (Ordinate axis in Fig. 1C).

The highly admixed populations with higher diversity in haplotype frequency may be challenging to properly classify, as can be seen from the correlation between the admixture error ‖ *E*_*i*_ ‖ and the admixture entropy *Ent*(*A*_*i*_) (Fig. 1C).

Note that the observed correlation may be the result of a cofactor such as the population size. To neutralize such an effect a partial correlation was performed (i.e. a correlation over the residual of the regression on the population size), with a Spearman partial correlation of 0.4 (p<1.e-4.). Interesting correlations also emerge between the entropy of the haplotype frequencies and the error, with a low error for high entropy (more uniform distribution of haplotypes). Moreover, as could be expected size is correlated with a lower error and a lower entropy of A, representing the bias toward the more precise representation of larger populations (Supp Mat. Fig. S2).

The admixture results (Fig. 1B) are consistent with the phylogenetic analysis based on the *Fst* distance between the HLA haplotype frequencies of the observed populations (Fig. 2A), where leaves are colored per *PP*(*i*). The tree splits into primary ethnic groups. Similar populations from different registries (e.g. populations of African ancestry from the Australian and US registries) are neighbors in the lineage tree and have similar *PP*(*i*) values, showing that technical differences between the methods used by different registries have a limited effect on the resulting admixture. The trees based on HLA alone effectively reproduce most details of current known population admixture, including the fine details of Asian and Jewish populations, as well as the admixture of Latin American populations[56, 57].

**Figure 2:**
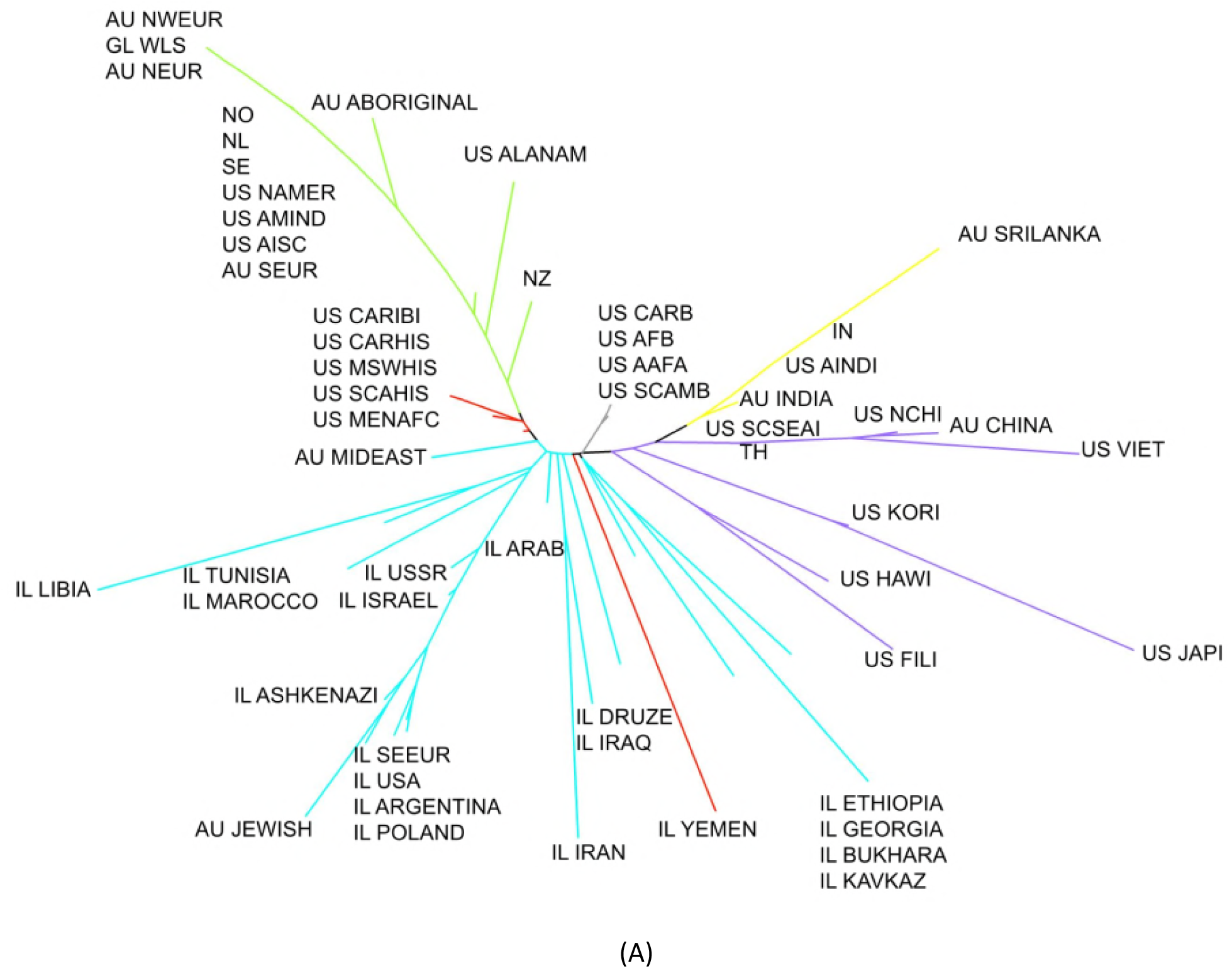
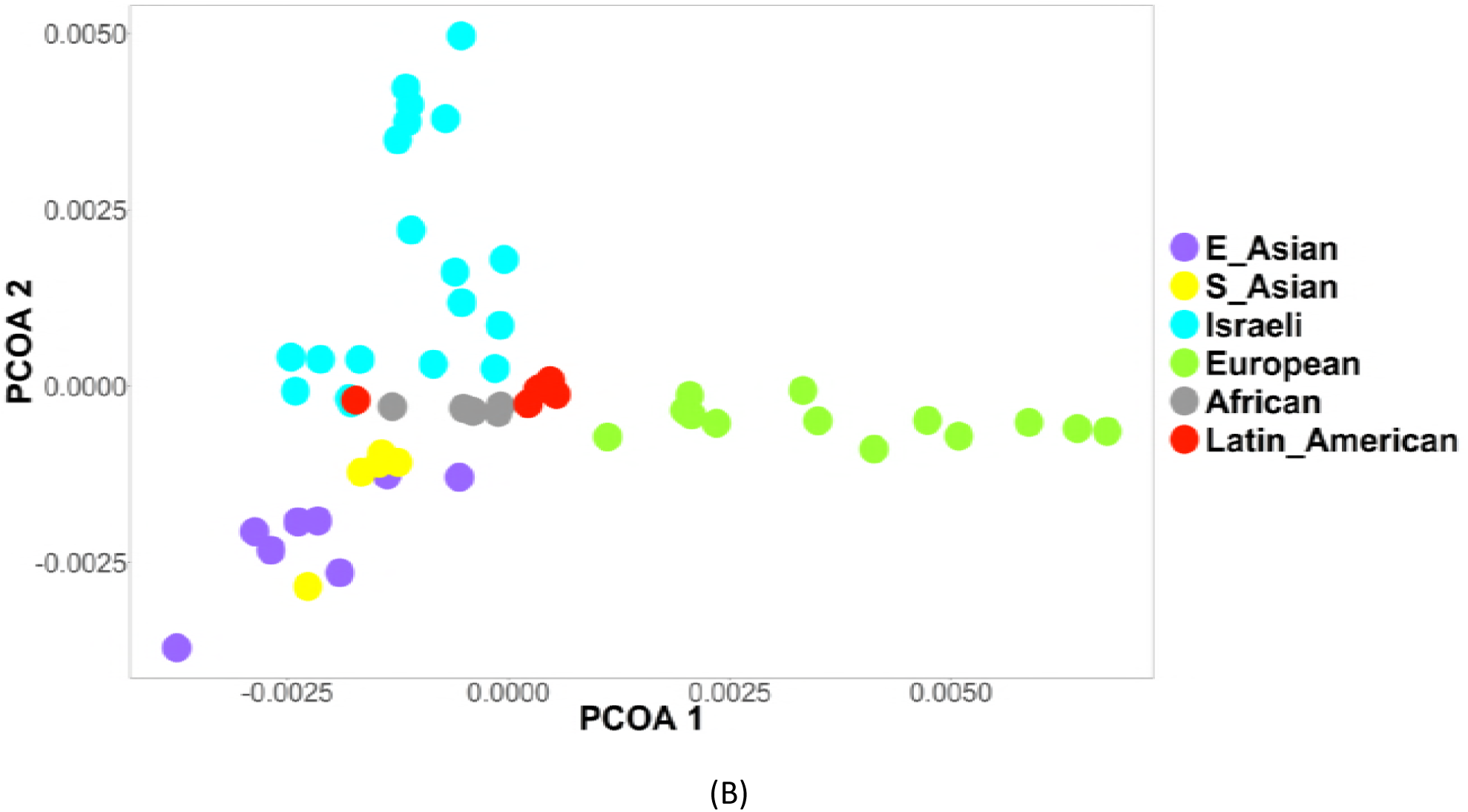
A) Unrooted neighbor-joining (NJ) phylogenetic trees based on the Fst matrix calculated from five-locus haplotype frequencies of the 56 studied populations. The tree is divided into branches delineating the main ethnic groups in the current analysis: Jewish, African, European, Hispanic, East and South Asians. Color coding shows concordance between the estimated admixture and tree branches. The only exceptions are highly admixed populations, such as Ethiopian Jews. B) PCOA (Principal Coordinate Analysis) analyses of the distances as defined by pairwise population Fst. The population have been grouped into broad regions: South Asian, East Asian, Pacific Islanders, Jews, European, African and Hispanic. More plots are provided in supplementary materials.

Some interesting trends in the tree, beyond reproducing the main human lineages, are the proximity between American Indians and European populations, which is previously reported[58, 59]. This can be seen also by the significant European OP component of Native American populations (Fig. 1B). The same trend holds even more for Australian Aboriginal populations which agrees with the high admixture between these groups and settlers reported in the literature[59-61]. Israeli Arabs and Druze are closer to their Jewish neighbors than to Middle Eastern or North African populations in the US or Australia. This is probably the result of the broad classification of these groups in the US and Australian registries[56].

The classification of populations based on their leading OP agrees with PCOA analyses of the pairwise population Fst distances based only on HLA. The PCOA is also in agreement with the geographical spread of the populations. The first PC (horizontal axis in Fig. 2A) represents an East-West division of populations, separating the European from all other populations. The second PC (vertical axis in Fig. 2A) separates Middle Eastern and Israeli groups from Asian populations. The African populations are in the center of the PCOA and the phylogenetic tree as expected from out of Africa evolutionary theories of human ancestry[62, 63]. Note the agreement of *PP*(*i*) with the PCOA map, further showing that the admixture captures the main axes of variance in the human genetic development (Other dimensions of the PCOA are given in Supp. Mat. Fig. S2).

To compare our results to existing methods, we performed an MCMC based admixture analysis on a subset of genome-wide ancestry informative marker SNPs in the 1000 genome (1KG) populations (Fig. 3A) and compared the result to a subset of geographically similar populations in our study where admixture was estimated using the NMF-based analysis and HLA haplotype frequencies. As seen in Fig. 3A, significant similarities can be seen between the NMF-based (top) and MCMC estimated admixtures (bottom), especially in Chinese, Japanese, African and European populations. We observed more splits in the Latin-American 1KG populations (MXL, PUR, CLM) in the MCMC admixture. This could be the result of over representation of these samples in the 1KG project compared with the registry samples. Note that the registry data does not contain a specific Iberian Spanish population, and the European admixture in registry Latin American populations is usually less than Iberian populations because of the Amerindian influence.

**Figure 3.**
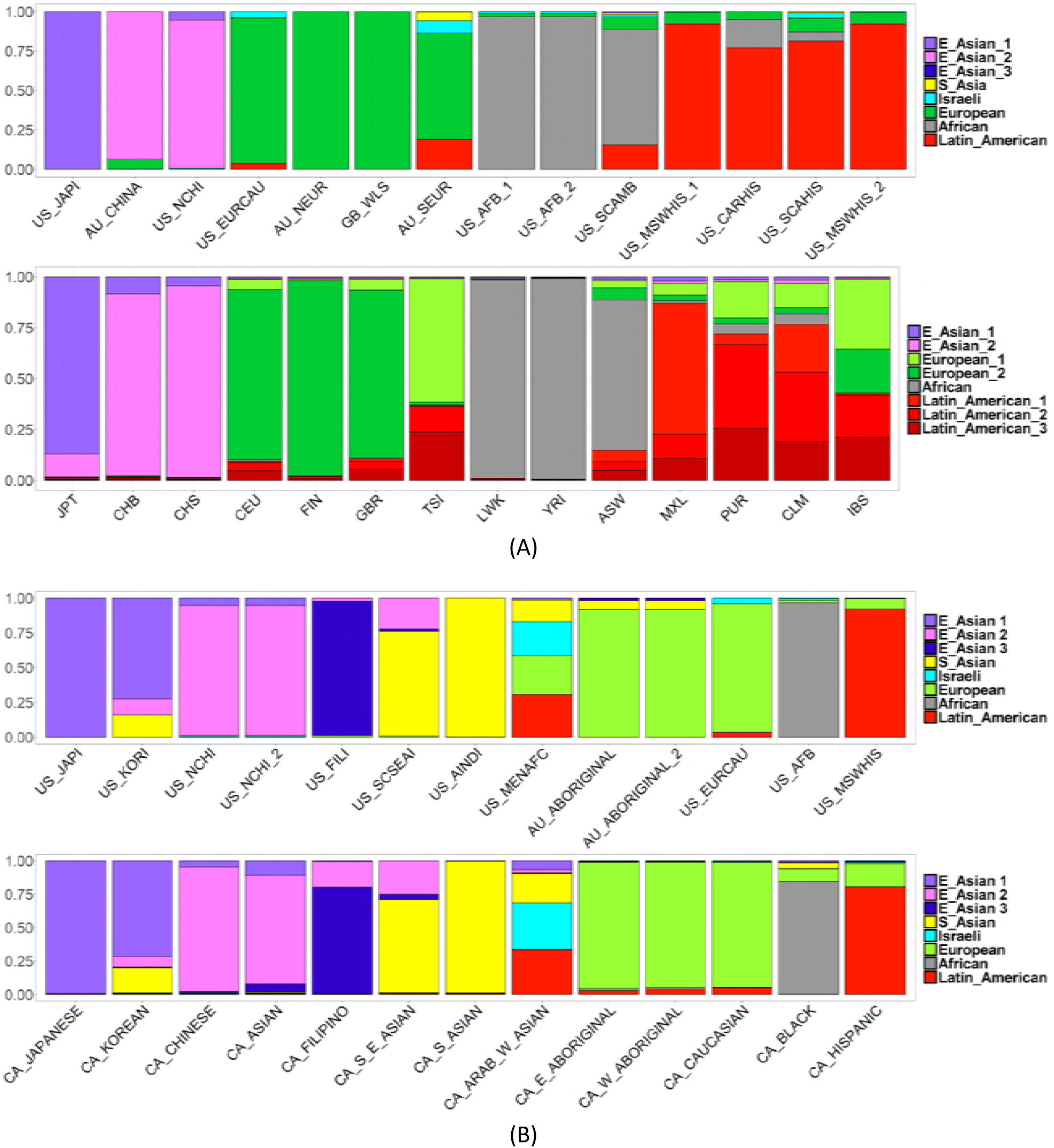
A) Comparison between admixture on 1,000 genome data from STRCUTURE analysis (lower plot) and from the current NMF (Non-negative Matrix Factorization) based analysis (upper plot). The colors represent the OPs as produced in STRCUTURE and the NMF. For each population in the 1K Genome data, the most similar population in the registry was used. B) Validation bar plot of Canadian population admixture based on the OPs (Original Populations) producing the admixture in Figure 1B. All haplotype frequencies were reduced to four-locus resolution like genotypes provided from the Canadian OneMatch registry.

One application of the presented methods is the detection of the composition of untested groups to understand their sub-structure. To estimate the composition of unknown populations, we performed an admixture analysis using the estimated OPs to compute admixture in groups from the Canadian OneMatch registry, without the limit that the entire composition should be explained by a combination of the OPs:

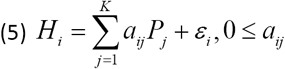

HLA haplotype frequencies from the Canadian registry were estimated at a four-locus resolution. To estimate the admixture of the Canadian populations, we dropped the resolution of the OPs haplotype frequency distribution to four-locus by summing over all five-locus haplotypes that differ only in HLA-DQB1. We then computed the admixture of the new observed populations using Eq. (5).

For most Canadian populations, the estimated admixture via OPs agreed with self-reported race and ethnicities and with similar groups from other registries in the US, Australia and Europe (Figure 3B). The similarity between the US and Canadian admixtures is clearly demonstrated in the admixture estimation of the Canadian Asian, African, European, Latin American, Korean, and Middle Eastern populations. However, some populations, such as Aboriginal and Filipino Canadians show a slightly different composition. For example, the Canadian Aboriginal population is composed not only of the Australian Aboriginal population components, but also has fragments from African and Hispanic populations. This could be attributed to different patterns of population migration and historical events.

## Discussion

We present an algorithm that dissects population genetic admixture, based on HLA haplotype frequencies, into OP components in the presence of background LD and high polymorphism. While traditional admixture and lineage models typically rely on many biallelic genome-wide loci, we demonstrate that HLA is polymorphic enough to allow for a clear delineation of population composition using a single genomic region. The admixture problem, shown equivalent to a Non-engative Matrix Factorization analysis, is more computationally efficient than traditional admixture models and allows for the admixture of a large number of populations with an extensive number of haplotypes (in the order of a million haplotypes). To our knowledge, this is the first algorithm to dissect population admixture with specific focus on HLA and a validation framework using a dataset with the presented magnitude.

We developed and applied our method to the haplotype frequencies of 56 populations from different adult volunteer stem-cell registries representing over 3.5 million donors. We showed that the resulting admixture is consistent with the known ethnic composition, recent history and SNP based admixture.

The results of the admixture and phylogenetic analyses show a clear distinction between Asian, African, European and Israeli populations. Expectedly, the Latin-American and Middle Eastern Arab populations were more admixed than other populations and contained a notable European component. Our method was also able to distinguish East Asian from south Asian and Pacific Islander populations such as New Zealand. Some of the analyzed populations were more admixed than others; for example, most East Asian populations had a single predominant OP while Caribbean and Middle Eastern groups were more admixed. A distinct difference emerged between Israeli and non-Israeli populations. Within the Jewish Israeli populations, phylogenetic analysis separated several distinct groups: Ashkenazi, North African, Central Asian, and Yemenite and Ethiopian Jews. Interestingly, the Jewish populations were mainly admixed with Caucasian and south Asian populations, probably representing historic migration and modern admixture events. Our results showed over 50% European admixture in Native American and Australian aboriginal populations, suggesting large recent admixtures (Figure 1B). Additionally, we have reported in a previous study a degree of over reporting of self-identified Native-American race among Be The Match donors [64] that did not completely coincide with the genetic composition of reporting individuals, some of whom were found to have substantial European admixtures. Our phylogenetic analysis showed comparable results (Figure 2A). The general division of branches agreed with previous phylogenies of general human populations

A limitation of the current methods is its restriction to phased HLA haplotype frequencies. The current implementation cannot be applied to SNP or other unphased marker data. Additionally, the presented admixture assignments are estimated at the population level, i.e. each haplotype is assigned a most likely OP, but a relative fraction of OP cannot be assigned to an individual. Thus, the presented method is complimentary to existing unphased admixture models.

Beyond the theoretical importance of NMF based admixture methods, the presented results are useful in the context of hematopoietic stem cell transplantation (HSCT). Current methods to estimate haplotype frequencies and detect optimally matched HLA donors are based on Self-Identified Race and Ethnicity (SIRE), with an assumption of endogamy within the same SIRE. The probability to find an optimally HLA matched stem-cell donor is higher within the same ethnic group of a searching patient [10]. However, both admixture and phylogenetic results presented here show that some SIRE groups are composed of multiple endogamous populations and some have similarities in haplotype frequency distribution. The presented algorithm can therefore help dissect these multiple SIRE groups into a multitude of sub-populations based on defined haplotype distributions, which would ultimately streamline the matching process for HSCT.

## Acknowledgement

We would like to thank the following registries for allowing their data to be used for this study: National Marrow Donor Program/Be The Match, USA, Ezer Mizion Bone Marrow Donor Registry, Israel, OneMatch Stem Cell and Marrow Network, Canada, Australian Bone Marrow Donor Registry, Australia, Matchis: the Dutch Centre for Stem Cell Donors, The Netherlands, Norwegian Bone Marrow Donor Registry, Norway, New Zealand Bone Marrow Donor Registry, New Zealand, Tobias Registry of Swedish Bone Marrow Donors, Sweden, Thai National Stem Cell Donor Registry, Thailand, Welsh Bone Marrow Donor Registry, Wales, United Kingdom.

The authors affirm that all data necessary for confirming the conclusions of the article are present within the article, figures, and tables or in published datasets.

This work was supported by a grant from the Department of the Navy, Office of Naval Research (N00014-16-1-2020).

